# Regional similarity to male-typical brain architecture links childhood ADHD to neurophysiological aging in later adulthood

**DOI:** 10.64898/2026.09.27.754764

**Authors:** Raluca Petrican, Alex Fornito, Amber John, Sophie Pendered

## Abstract

Identifying early life biomarkers of suboptimal aging trajectories could help design interventions to delay or even prevent the onset of pathological conditions in older adulthood. To address this issue, we capitalize on evidence that the prevalence rates and presentation of neurodevelopmental and neurodegenerative conditions tend to vary between males and females, thereby raising the possibility that processes involved in sexual differentiation could help connect brain maturation and aging outcomes. Here, we focus on attention deficit hyperactivity disorder (ADHD), a neurodevelopmental condition with male-biased prevalence, and examine the relevance of its genetic risk and functional brain profile to aging outcomes. We analyzed multimodal data from a child sample enriched in ADHD cases and a neurologically intact adult sample. Among 6-to-12-year-olds (N =522), genetic vulnerability to pathological aging, indexed via polygenic risk scores (PRSs) for Alzheimer’s Disease (AD), predicted ADHD diagnosis across sexes, regardless of psychiatric history and PRSs for multiple disorders (including ADHD). Irrespective of age, boys and girls diagnosed with ADHD showed preferential alignment with male-, rather than female-, typical functional connectivity patterns in metabolically active transmodal areas which are enriched in 5-HT_2,4_, D1, and mGLU5 receptors. In an independent sample aged 36-90 years (N = 345), the ADHD brain profile correlated with higher levels of testosterone, physiological markers of reproductive aging and poorer performance on AD-relevant cognitive tasks among older individuals of either sex. Our results imply that neurodevelopmental conditions could provide insights into vulnerability to suboptimal aging trajectories and could help guide early interventions targeting at-risk individuals.

---

Recent years have seen increasing commitment to personalizing neuropsychiatric detection and intervention paradigms through the adoption of a life course approach to mental and physical health (Lopez et al., 2021; van Zwieten et al., 2025). This perspective is rooted in mounting evidence that pathological aging conditions have an extended, often decades long, phase marked by subclinical and preclinical changes (De Strooper & Karran, 2016; Hampel et al., 2023), during which interventions could slow down or even prevent progression towards clinical onset.

Here, we took a first step towards characterizing early life biomarkers of suboptimal lifespan developmental trajectories. Specifically, capitalizing on evidence of robust sex differences in maturation and aging (e.g., (Bachmann et al., 2023; Barth et al., 2015; Keller et al., 2025; Lopez-Lee et al., 2024; Ma et al., 2024; Newhouse & Dumas, 2015), we reasoned that overlapping neurobiological mechanisms related to sex differences in brain function could relate to both developmental challenges in early life and cognitive-affective outcomes in later adulthood. Alignment with sex-typical functional brain architecture has been recently linked to adolescent risk for psychiatric conditions with sex-biased prevalence. Specifically, across males and females alike, increasing alignment with female-typical, rather than male-typical, FC patterns from sensorimotor to transmodal brain regions has been connected to individual differences in depression and anxiety, whereas increasing alignment with male-typical, rather than female-typical, FC patterns along the same axis has been related to behavioral problems (Petrican et al., 2025). These findings raise the possibility that regional variability in relative alignment with sex-typical brain architecture could help link neurodevelopmental conditions with sex-biased prevalence to patterns of neurocognitive aging.

To address this issue, we focused on attention deficit hyperactivity disorder (ADHD), one of the most widely diagnosed neurodevelopmental conditions, which shows male-biased prevalence (Martin et al., 2024) and is currently thought to affect around 8% of children and adolescents worldwide (Ayano et al., 2023). Challenges in modulating cognition and behavior based on environmental demands are common in ADHD and may underpin many of its sequelae, including lower educational attainment and poorer occupational outcomes (Barkley, 1997; Castellanos & Tannock, 2002; Thapar & Cooper, 2016; Willcutt et al., 2005). Nonetheless, the heterogenous clinical profile of ADHD and its wide range of strengths and difficulties stem from complex interactions among mental functions which partly overlap with those most vulnerable to aging, such as cognitive control (e.g., inhibition, working memory) and memory storage (Frisoni et al., 2022; Mirabella, 2021; Thapar & Cooper, 2016). Beyond cognition, ADHD has further links to suboptimal aging trajectories via immune and cardiometabolic dysregulation. For instance, a three-generation population- based cohort study linked genetic risk for ADHD to greater peripheral inflammation, and incidence of both metabolic and cardiovascular disorders (Du Rietz et al., 2024).

To our knowledge, the neurobiological profile of childhood ADHD is yet to be interrogated in relation to aging outcomes. Here, we probed this issue through three interconnected lines of inquiry. First, we investigated whether childhood ADHD is related to genetic risk factors predictive of suboptimal aging outcomes, specifically, dementia risk, and whether this relationship is independent of sex, age, psychiatric history and genetic vulnerability to other mental health conditions. Second, across sexes, we probed whether regional patterns of alignment with female- vs male-, typical brain architecture, which are associated with childhood ADHD, would overlap with those linked to poorer functioning in middle to older adulthood, specifically, steeper physiological decline, as well as reduced episodic memory and cognitive control (Ourry et al., 2024; Xiong et al., 2025; You et al., 2024). A robust overlap would raise the possibility that patterns of regional alignment with sex-typical brain architecture may be one of the age-invariant neural correlates of episodic memory and cognitive control, which could shed some light on why children diagnosed with ADHD may be at risk for suboptimal aging outcomes. The later life physiological decline measure indexed immune/metabolic dysregulation, which have been related to ADHD (Du Rietz et al., 2024) together with gonadal hormone levels, which play a key role in normative neural sexual differentiation (Gegenhuber et al., 2022; Küchenhoff et al., 2024), as well as in neurocognitive aging (e.g., (Barth et al., 2015; Lopez-Lee et al., 2024; Newhouse & Dumas, 2015)). Given its male-biased prevalence (Martin et al., 2024), we expected that childhood ADHD would correlate most robustly with regional patterns of alignment with male-, rather than female-, typical FC and, in the adult sample, also relatively greater testosterone levels (after controlling for sex) (Gegenhuber et al., 2022; Küchenhoff et al., 2024). We expected for the ADHD-brain effects to hold across boys and girls because our underlying assumption was that the greater prevalence of ADHD among male individuals is partly accounted by the fact that the key neural features of ADHD, which are observed across sexes, tend to resemble more closely a typically male, rather than female, brain architecture. This hypothesis was directly tested in control analyses corresponding to the planned Analyses 2 and 3.

Finally, we probed the neurochemical correlates of the ADHD functional brain profile, seeking to identify the neurotransmitter systems likely to link neurodevelopmental conditions to aging outcomes. This analysis included the neurotransmitter systems implicated in both ADHD and pathological aging, specifically, the dopaminergic (DA), glutamatergic [GLU]/GABA-ergic and noradrenergic systems (Koirala et al., 2024; Parr et al., 2025; Pertermann et al., 2019; Pilotto et al., 2025; Soares et al., 2024; Thapar & Cooper, 2016; Tsikonofilos et al., 2025), as well as those more broadly relevant to attention and cognitive control (i.e., cholinergic [Ach] (Ananth et al., 2023; Ballinger et al., 2016)) or psychopathology (i.e., serotonin [5-HT], (da Cunha-Bang & Knudsen, 2021; Erritzoe et al., 2023; Svensson et al., 2021).

## 2. Methods

The methods are summarized below and detailed in the Supplemental Information (SI). The selection criteria for the target groups are outlined in Figure 1.

**Figure 1.**
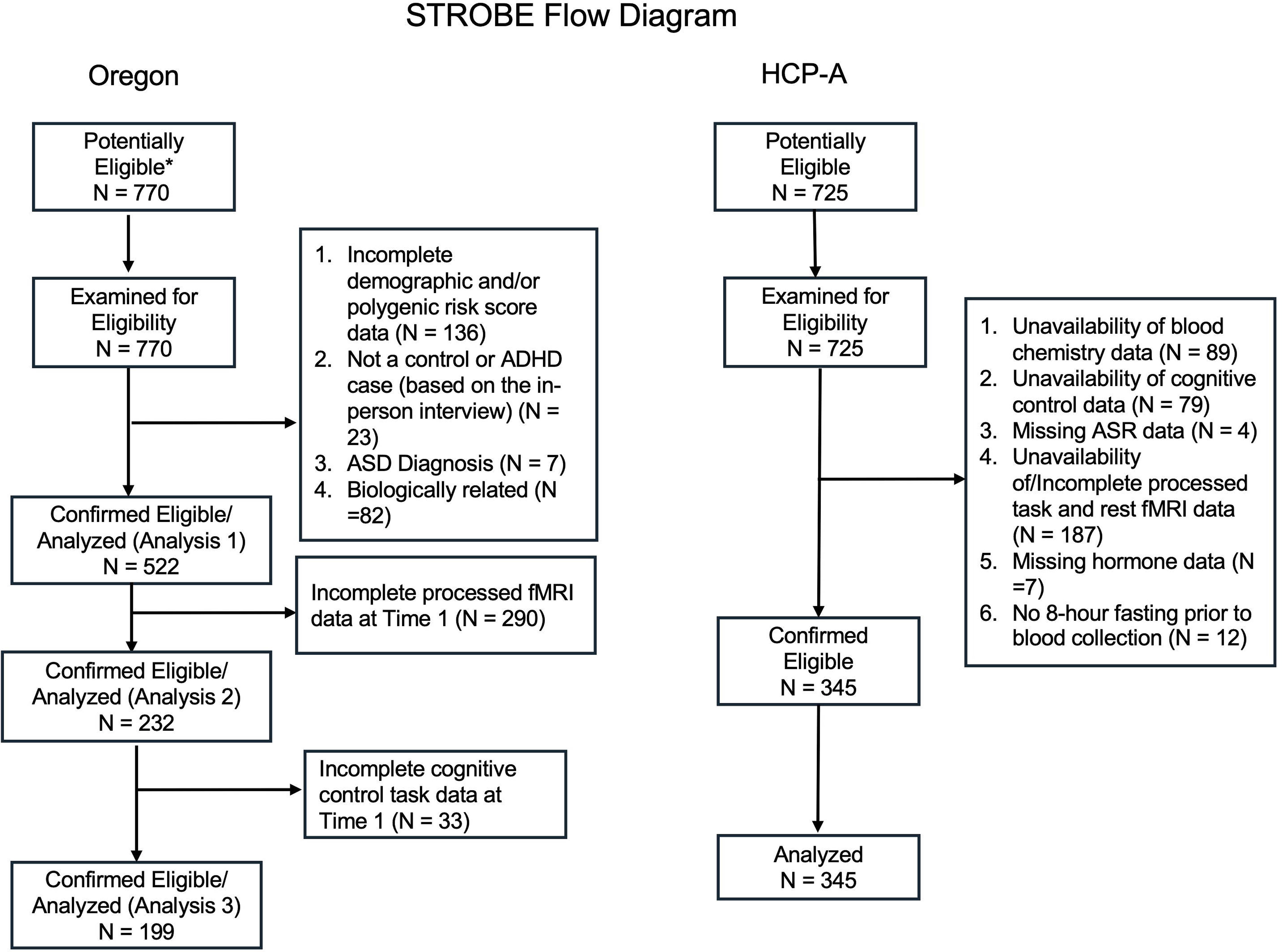
STROBE flowchart describing sample selection for the Oregon ADHD 1000 and HCP-A cohorts.

### 2.1 Participants

We analyzed data from 522 non-sibling children (196 girls; age range: 6-12 years) who participated in the Oregon ADHD-1000 study (Nigg et al., 2023) and 345 unrelated participants (195 women, age range: 36-90 years) from the Human Connectome-Aging (HCP-A) (Bookheimer et al., 2019). Most participants were White, predominantly right-handed and none self-reported being pregnant.

### 2.2 Psychiatric History and Genetic Risk for Psychiatric Disorders

#### Oregon

Lifetime history of major depressive disorder (MDD), anxiety (i.e., generalized anxiety disorder [GAD], social anxiety, separation anxiety) and oppositional defiant disorder (ODD) was derived from parental responses on the Kiddie Schedule for Affective Disorders and Schizophrenia (KSADS) (Kaufman et al., 1997) at the first study visit. Polygenic risk scores (PRSs) for sporadic Alzheimer’s Disease (AD), the most common cause of dementia over the age of 60 ((ADI), 2025) ADHD, MDD, anxiety (ANX), schizophrenia (SCZ) and bipolar disorder (BD) were computed using the summary statistics from large case-control genome-wide association studies (GWASs) focused on each condition (Demontis et al., 2023; Kunkle et al., 2019; Major Depressive Disorder Working Group of the Psychiatric Genomics Consortium. Electronic address & Major Depressive Disorder Working Group of the Psychiatric Genomics, 2025; O’Connell et al., 2025; Strom et al., 2026; Trubetskoy et al., 2022). For AD, we computed separate PRSs based on APOE vs no-APOE risk alleles, based on evidence that the two relate to distinguishable neurocognitive trajectories (i.e., memory-focused for the former vs cognitive control-focused for the latter, (Frisoni et al., 2022)). PRSs for behavioural disorders (e.g., conduct disorder, ODD) were not included because we could not locate any pertinent case-control GWASs. Disorder-specific PRSs were computed with the PLINK genetic analysis toolset (Chang et al., 2015) as the sum of the corresponding risk alleles weighted by their associated posterior effect size estimates which were outputted by PRS-CS across its standard range of global shrinkage values (i.e., phi=1e-6, 1e-4, 1e-2, 1) (Ge et al., 2019). Because we had no reason to favour a specific *phi* value, the disorder-specific PRSs were averaged across the four scrutinized *phi*’s (for each disorder, PRS intercorrelations ranged from .42 to .99). The CCA results based on individual phi values are presented in Figure 2.

**Figure 2.**
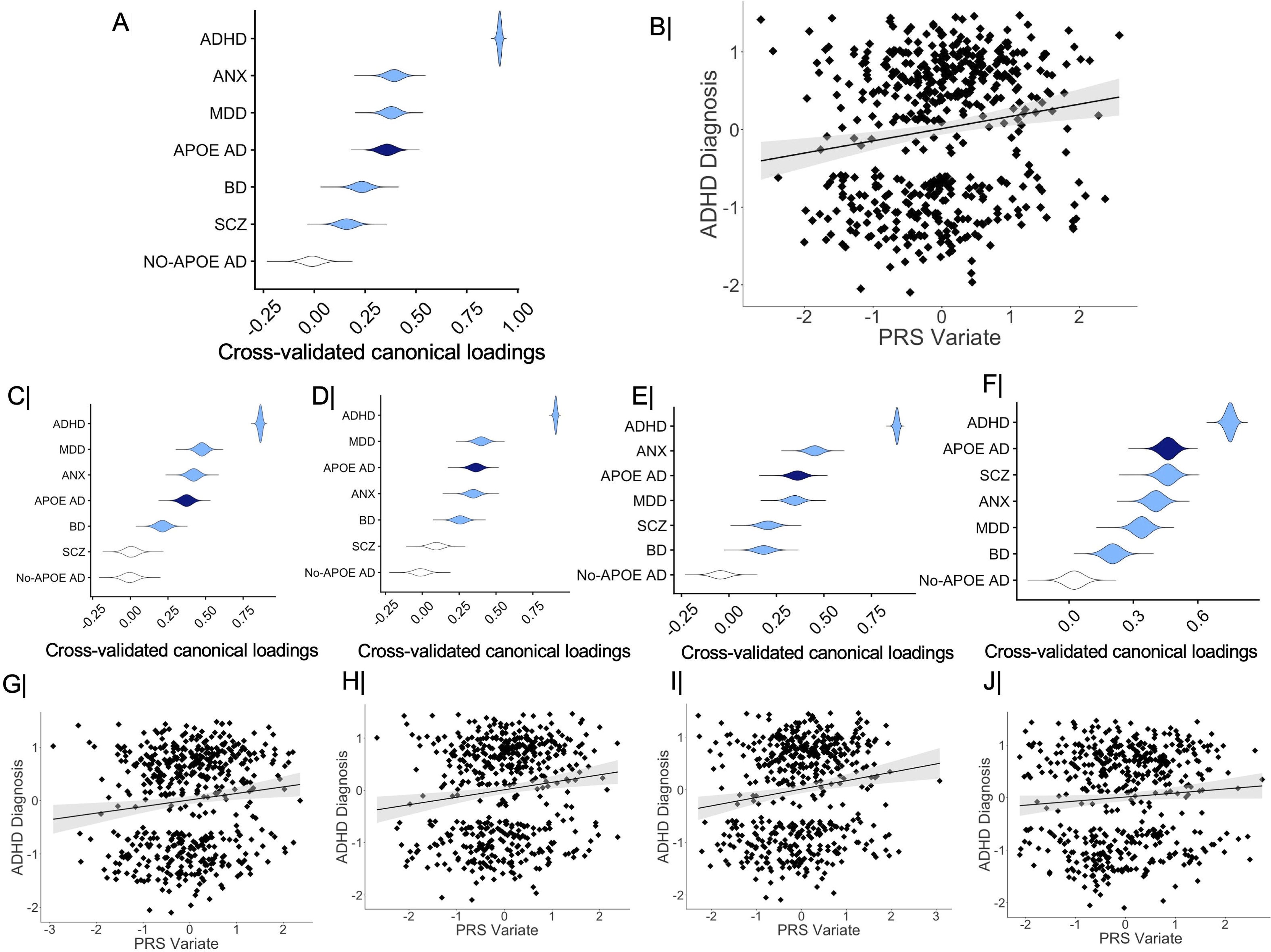
Polygenic risk scores (PRSs) linked by CCA to clinically diagnosed ADHD in the Oregon sample. The partial correlation coefficients describing the relationship between the measured polygenic risk scores and the predicted value of their corresponding canonical variate across all test CCA folds are presented in panels A (disorder-specific PRSs averaged across all phi-values), C (disorder-specific PRSs for a phi-value of 1), D (disorder-specific PRSs for a phi-value of 1e2), E (disorder-specific PRSs for a phi-value of 1e4) and F (disorder-specific PRSs for a phi-value of 1e6). The violin plots in panels A, C-F depict the distribution of partial correlation coefficients across the 100,000 bootstrap samples from the cross-validation procedure. Panels B and G-J contain the scatter plot describing the linear relationship between a diagnosis of ADHD and the CCA variate pictured in panels A, C-F. In panels A and C-F, gray colored violin plots indicate non-robust canonical loadings (based on the bootstrapped 99% CIs from the cross-validation procedure). CCA = canonical correlation analysis. CI = confidence interval. ADHD = attention deficit hyperactivity disorder. Anx = anxiety. MDD = major depressive disorder. SCZ = schizophrenia; BD = bipolar disorder. AD = Alzheimer’s Disease. PRS = polygenic risk score.

The discovery and, therefore, the linkage disequilibrium (LD) reference from 1000 Genomes populations were of European descent. To account for population stratification effects related to ancestry, the first three genomic principal components (PCs) (eigenvalues > 2, eigenvalues for the remaining PCs < 1.3) extracted with the PC-AiR function from the Bioconductor package GENESIS (Gogarten et al., 2019) were covaried out in all the analyses featuring the PRSs (see also (Mooney et al., 2024).

#### HCP-A

Psychological functioning, specifically, self-reported symptoms of ADHD, depression, anxiety, antisocial personality, avoidant personality and somatic disorder, were measured with the Adult Self-Report Scale (ASR, (Achenbach, 2009).

### 2.3 Cognitive Functioning

#### Oregon

Cognitive differences relevant to the clinical profile of childhood ADHD were assessed in the fixed order outlined below after a washout of stimulant medications of at least 7 half-lives (use of other medications was an exclusion criterion) (Nigg et al., 2023).

The children thus completed measures of attentional control (i.e., Go/Stop task), learning (i.e., spatial span [forward, backward] and n-back working memory tasks) and cognitive flexibility (i.e., task switching).

#### HCP-A

We analyzed measures of attentional control (i.e., inhibition [Flanker task]), learning (i.e., working memory [list sorting]) and cognitive flexibility (dimensional card sorting task) relevant to ADHD. Cognitive deficits relevant to suboptimal aging, specifically, AD risk, were gauged via an episodic memory (Picture Sequence Task) and a processing speed task (Pattern Comparison Task). The measures have been described in detail elsewhere (Bookheimer et al., 2019)).

### 2.4 Physiological Aging (HCP-A)

#### PhenoAge

Physiological wear-and-tear, an indicator of suboptimal aging and a putative risk factor for AD (Irwin & Vitiello, 2019), was estimated with PhenoAge, a well-validated algorithm for predicting health and mortality through blood chemistry indicators of immune and metabolic system integrity (Levine, 2013). Individuals with a more advanced PhenoAge (relative to their chronological age) show greater physiological wear-and-tear and, thus, are at greater than expected (by chronological age) risk for disease and mortality (Belsky et al., 2015; Levine, 2013). The biological age prediction model was trained and validated in two samples (aged 20-90 years) from the National Health and Nutrition Examination Survey (NHANES) (https://wwwn.cdc.gov/nchs/nhanes/Default.aspx). For each HCP-A participant we computed the difference between the age estimated by the PhenoAge algorithm and their chronological age. Positive values indicated premature, whereas negative values indicated delayed, aging.

#### Gonadal hormones

Levels of estradiol (E2), testosterone, luteinizing hormone (LH) and follicle stimulating hormone (FSH) were extracted from blood samples following confirmed fasting for 8 hours prior to testing. To account for variable hormone, especially during perimenopause, blood samples for women aged 45-55 years were collected during days 2-6 of their menstrual cycle (Bookheimer et al., 2019). All the analyses that included gonadal hormones controlled for the participant’s sex.

### 2.5 fMRI Data Acquisition and Processing

The functional neuroimaging data from all three samples had been processed with the HCP Preprocessing Pipelines (Glasser et al., 2013), which were enhanced for the Oregon sample with additional corrections for in-scanner head motion (Nigg et al., 2023) (see Figure S7 for a summary of motion-functional connectivity associations in the Oregon sample). The acquisition and preprocessing protocols had been already published (Harms et al., 2018; Nigg et al., 2023; Smith et al., 2013) and are summarized in the SI.

### 2.6 Sex-Related FC Strength Gradient

Sex-differential FC strength patterns were characterized among the biologically unrelated healthy young adults (N = 336, aged 22-30 years) from the Human Connectome Project (HCP) who had reached reproductive maturity and were thus expected to show maximal distinguishability in brain function in accordance with their biological sex (see {Petrican et al., 2025 and SI for details on the computation of the sex-related FC strength gradient in the HCP sample). Specifically, we regressed each of the 44850 parcel-to-parcel FC indices against sex and a number of variables which have been related to FC strength and on which complete matching between sexes is not feasible, specifically, age, income, employment status, race, handedness, average in-scanner motion, intelligence and psychopathology. For each parcel-to-parcel FC index, the *t*-statistic corresponding to “sex” was taken as a measure of the robustness with which the respective FC index differentiates between men and women. This decision was made in order to avoid prioritising FC indices that show large sex differences in a subsample, but not the entire HCP group. Nonetheless, in line with the intentional homogeneity of the selected HCP group, the *t*-statistic and corresponding beta weight value associated with “sex” for a given parcel-to-parcel FC index were very strongly correlated (median *r* of .96, range: .79-.995).

### 2.7 Alignment with the HCP sex-related FC gradient in the Oregon and HCP-A samples

For each Oregon and HCP-A participant, relative alignment with female- vs male-typical FC patterns was estimated as rank order correlation-based similarity to the sex-related FC strength gradient from the HCP sample (i.e., the t-statistic matrix associated with the effect of sex on FC strength). We relied on Spearman’s, rather than Pearson’s, correlations because we had no reason to assume a linear relationship between the HCP *t*-statistics indexing the reliability of sex differences in each parcel-to-parcel FC and the corresponding parcel-to-parcel FC strength from an Oregon or HCP-A participant. Use of Spearman’s correlations also safeguarded against the skewed distribution of some regional FC profiles from the Oregon or HCP-A samples (Riffenburgh & Gillen, 2020). A positive correlation coefficient reflects greater similarity to the male-, rather than female-, typical FC strength profile. A negative correlation coefficient indicates higher similarity to the female-, rather than male-, typical FC strength profile. Because the HCP *t*-statistic matrix is based only on young adult FC profiles, anchored at one end in female prototypicality and, at the other, in male prototypicality, both positive and negative rank order correlation coefficients indicate similarity to a young adult brain profile. Figure 3-A, B contains a schematic representation of the sex-related FC gradient analyses.

**Figure 3.**
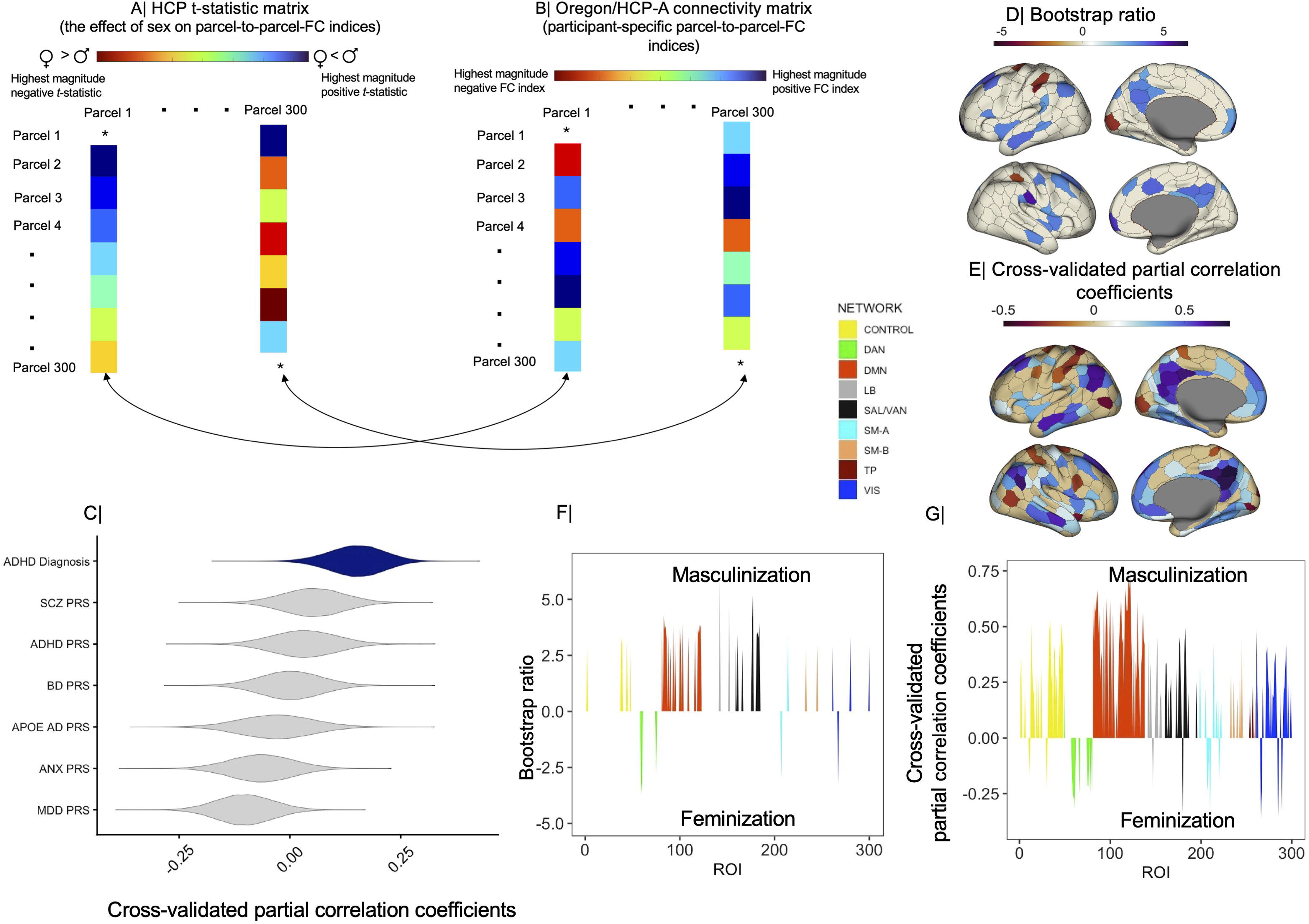
The brain LV from the behavioral-PLS analysis linking a diagnosis of ADHD and its PRS correlates from Analysis 1 to alignment with sex-typical brain architecture in the Oregon sample. Panels A-B depict the analytic steps through which alignment with female- vs male-typical FC patterns was estimated. For each Oregon or HCP-A participant, relative alignment with female- vs male-typical FC patterns was computed as the rank-order correlation coefficient between each column of their own FC matrix (i.e., each parcel’s FC with each of the remaining parcels in the atlas) (panel B) and the corresponding column in the HCP *t*-statistic matrix (panel A). A positive correlation coefficient indicates that the regional FC strength profile from an Oregon or HCP-A participant aligns with the ascending female-to-male FC strength gradient from the HCP (a more masculine FC strength profile). A negative correlation coefficient indicates that the regional FC strength profile from an Oregon or HCP-A participant aligns with the ascending male-to-female FC strength gradient from the HCP (a more feminine FC strength profile). *T*-statistic values corresponding to self-connections (panel A) or FC indices corresponding to self-connections (panel B) were set to zero. Panel C shows the correlation of the ADHD diagnosis and its PRS correlates from Analysis 1 with the brain LV scores in the cross-validated PLS analysis. The violin plots in panel A depict the distribution of these correlation coefficients across the 100,000 bootstrap samples from the discovery or cross-validated PLS analysis. Panel D depicts the ROI-specific weights/loadings on the brain LV identified with the discovery PLS analysis with a bootstrap ratio greater than 2.75 in absolute value (equivalent to a 99% CI). Panel F presents the same information aggregated by the Schaefer atlas networks. Panel E depicts the Schaefer ROIs robustly correlated (based on cross-validated 99% confidence intervals) with the predicted value of the brain LV from the cross-validation procedure. These are partial correlations controlling for the confounders listed for Analysis 2 under Results. Panel G presents Schaefer network-based distributions of partial correlations summarizing the ROI-specific results from panel (E). In panels F and G, positive values indicate greater alignment with male-, rather than female-, typical FC patterns (“masculinization”), whereas negative values indicate stronger alignment with female-, rather than male-, typical FC (“feminization”) for a specific ROI. As stated in the main text, alignment with sex-typical architecture was estimated in reference to resting state connectivity data from the Human Connectome Project. BSR = bootstrap ratio. PLS = partial least squares. LV= latent variable. CI = confidence interval. Schaefer networks: TP= temporo-parietal. SAL-VAN = salience/ventral attention. LB = limbic. DMN = default mode. DAN = dorsal attention. SM-A = somatomotor-A. SM-B =somatomotor-B. VIS = visual.

#### Validation of the sex-related FC strength gradient

The HCP sex-related FC strength models were validated in the Oregon and the HCP-A samples by confirming that in both samples the male participants showed stronger alignment in FC with the HCP men, whereas the female participants showed stronger alignment in FC with the HCP women (see SI). The validation and all the results reported below were replicated with HCP sex-related FC strength models which further controlled for brain size and female menstrual cycle characteristics, variables that reportedly impact structural brain characteristics (Küchenhoff et al., 2024) (see SI). Our comparative analyses of the Oregon and HCP-A data are rooted in the implicit assumption of a persistent pattern of alignment with sex-typical FC profiles from childhood to older adulthood. This was confirmed in supplemental tests depicted in Figures S8-9, which show that alignment with sex-typical FC profiles emerges gradually from birth onwards, with young adults constituting the most appropriate reference group because they reached reproductive maturation and would thus show maximal differentiation in sex-related FC. This pattern of results dovetails with previous reports that sex differences in brain structure and function can be observed at birth but they become most salient once reproductive maturation is reached (Ahmed et al., 2008; Gilmore et al., 2018; Hines et al., 2015; Pfeifer & Allen, 2021; Piekarski et al., 2023; Sato et al., 2004).

### 2.8 Statistical Analysis

The analyses featured two multivariate data-driven techniques, canonical correlation analysis (CCA) and partial least squares correlation (henceforth abbreviated as PLS) analysis. Our reliance on CCA and PLS was justified by evidence that psychiatric conditions are best characterized in relation to intrinsically correlated functional systems spanning multiple anatomical brain regions and levels of analysis/modalities (Graham et al., 2021; Insel et al., 2010; Williams et al., 2024; Williams et al., 2016).

Most of the reported tests (Analyses 1, 3-5) were conducted with CCA because of its superior ability to maximize cross-set associations (Mihalik et al., 2022). However, in Analysis 2, which featured datasets containing highly correlated within-set variables, we favored PLS because it outperforms CCA on reliability and reproducibility in such circumstances (McIntosh & Misic, 2013; Mihalik et al., 2022). All the permutation- and bootstrap-based tests used 100,000 samples. To maximize interpretability of the model estimates, the discovery CCA and PLS analyses were conducted on data that had not been residualized for any confounders. To demonstrate the robustness of our results, the CCA and PLS cross-validation controlled for well-known nuisance variables, as specified for each analysis. To optimize model estimation, the training sets used to cross-validate the Oregon PLS and CCA models were stratified by diagnosis (i.e., ADHD), whereas in the neurotypical HCP-A sample the training sets were stratified by sex (because the hormonal variables entered in this analysis show marked sex differences).

## 3. Results

### 3.1 Analysis 1: Across sexes, childhood ADHD is linked to genetic risk for multiple psychiatric conditions and APOE-related AD

We started by characterizing the genetic risk profile of ADHD in the largest group of biologically unrelated Oregon youths with available data on all the relevant variables (N = 522; 329 cases [236 males]; 193 controls [90 males]). A CCA linked a diagnosis of ADHD to the PRSs predicting risk for ADHD, MDD, ANX, SCZ, BD, and APOE-related vs APOE- unrelated AD. The extracted mode was successfully cross-validated after controlling for the children’s sex, age, race, maternal education, psychiatric history and scores on the first three genomic PCs (*r*_CV_ = .16, 95% CI = [.07; .24], permutation-based *p* = 22 x 10^-5^, Figure 2-B).

The results indicated that, irrespective of sex, children diagnosed with ADHD show higher genetic risk scores for ADHD and also, albeit to a much lesser extent, for MDD, ANX, APOE-related AD, BD and SCZ (Figure 2-A). The near-zero correlations of the APOE-related AD PRS with the remaining PRSs (*r*s from −.01 to .03) speak to its unique contribution to the genetic profile of ADHD (intercorrelations among the remaining disorder-specific PRSs from .09 to .60).

### 3.2 Analysis 2: Regional alignment with male-typical FC covaries with childhood ADHD across sexes

Having shown that a childhood diagnosis of ADHD is linked to genetic risk for multiple neuropsychiatric disorders, we next examined the potential brain underpinnings of these effects. We therefore conducted a PLS analysis to identify regionally specific patterns of alignment with female- vs male-typical FC linked to ADHD and its PRS correlates across sexes (cf Analysis 1, Figure 2). Availability of baseline resting state data constrained this analysis to 232 youths (150 cases [101 males]; 82 controls [39 males]) of the 522 entered in Analysis 1.

The discovery PLS analysis identified one robust brain-behaviour latent variable (LV) pair (shared variance 46.25%, permutation-based *p* = 5 x 10^-5^) linking relative alignment with female- vs male-, typical FC to ADHD and four of its six PRS correlates identified in Analysis 1 (ADHD, ANX, APOE-related AD, SCZ). However, only the relationship of the brain LV with ADHD survived cross-validation tests which controlled for the children’s sex, age, race, maternal education, psychiatric history and average in-scanner motion (*r*_CV_ = .16, 95% CI = [.04; .28], permutation-based *p* = .019, Figure 3-C). Thus, irrespective of sex and genetic etiology, a clinical diagnosis of ADHD related to widespread alignment with male-typical architecture, particularly in default mode (DMN), salience-ventral attention (SAL- VAN), Control and visual (VIS) regions, but relatively stronger alignment with female-typical FC in DAN areas (Figure 3-, D-G).

#### Sex differences in the relationship between ADHD and alignment with male-typical FC patterns

Although our objective was to identify patterns of alignment with sex-typical FC predictive of ADHD across sexes, we nonetheless explored the possibility that the brain- ADHD relationship may vary between girls and boys. To this end, we re-ran Analysis 2 with boys and girls modelled as separate groups and a diagnosis of ADHD as the “behavioral” within-group variable (the group-PLS model that also featured the six PRSs from Analysis 2 as within-group variables could not be cross-validated). This analysis yielded a brain- diagnosis LV pair which was successfully cross-validated (*r*_cv_ = 21, 95% CI [.10; .34], permutation-based *p* = 12.6 x 10^-4^). The group-PLS brain LV was almost identical with the one linked to ADHD in Analysis 2 (*r* of .96). As such, unsurprisingly, the regression analysis predicting the group-PLS brain LV from a diagnosis of ADHD, sex, and the interaction of the two, while adjusting for relevant confounds (i.e., age, race, maternal education, psychiatric history, average in-scanner motion) revealed a non-significant interactive effect of sex and diagnosis. This result is consonant with the interpretation that stronger alignment with male-typical, rather than female-typical, FC is observed among both boys and girls with ADHD.

### 3.3 Analysis 3: Increased alignment with male-typical FC patterns predicts ADHD-relevant differences in attentional and memory performance

Analysis 2 linked childhood ADHD to regionally specific patterns of alignment with sex-typical brain architecture. The next question was whether the observed pattern of stronger alignment with male-typical FC would correlate with, and, thus, be a plausible candidate mechanism for explaining the attentional and learning challenges relevant to ADHD. A related question was whether a potential link between ADHD-relevant alignment with sex-typical brain architecture and cognitive challenges would be modulated by genetic vulnerability to ADHD and other psychiatric conditions (ANX, MDD, APOE-related AD, BD, SCZ) (cf Analysis 1).

To address this question, we conducted a CCA using the 199 youths (126 cases [88 males]; 73 controls [37 males]) of the 232 from Analysis 2 with complete cognitive task data. The cross-validation tests controlled for the children’s sex, age, race, maternal education, psychiatric history, average in-scanner motion and the first three genomic PCs. Across all test folds, the ADHD-linked profile of increased alignment with male-typical FC was associated with poorer inhibitory control (i.e., longer time to stop on the Stop Signal task) and learning (i.e., reduced accuracy in the 1-back condition of the working memory task and on the forward span task), *r*_CV_ = .27, 95% CI = [.14; .42], permutation-based *p* = 32 x 10^-5^ (Figure 4- B, C). The brain-cognition relationship was weaker among children who were at higher genetic risk for anxiety (Figure 4-A).

**Figure 4.**
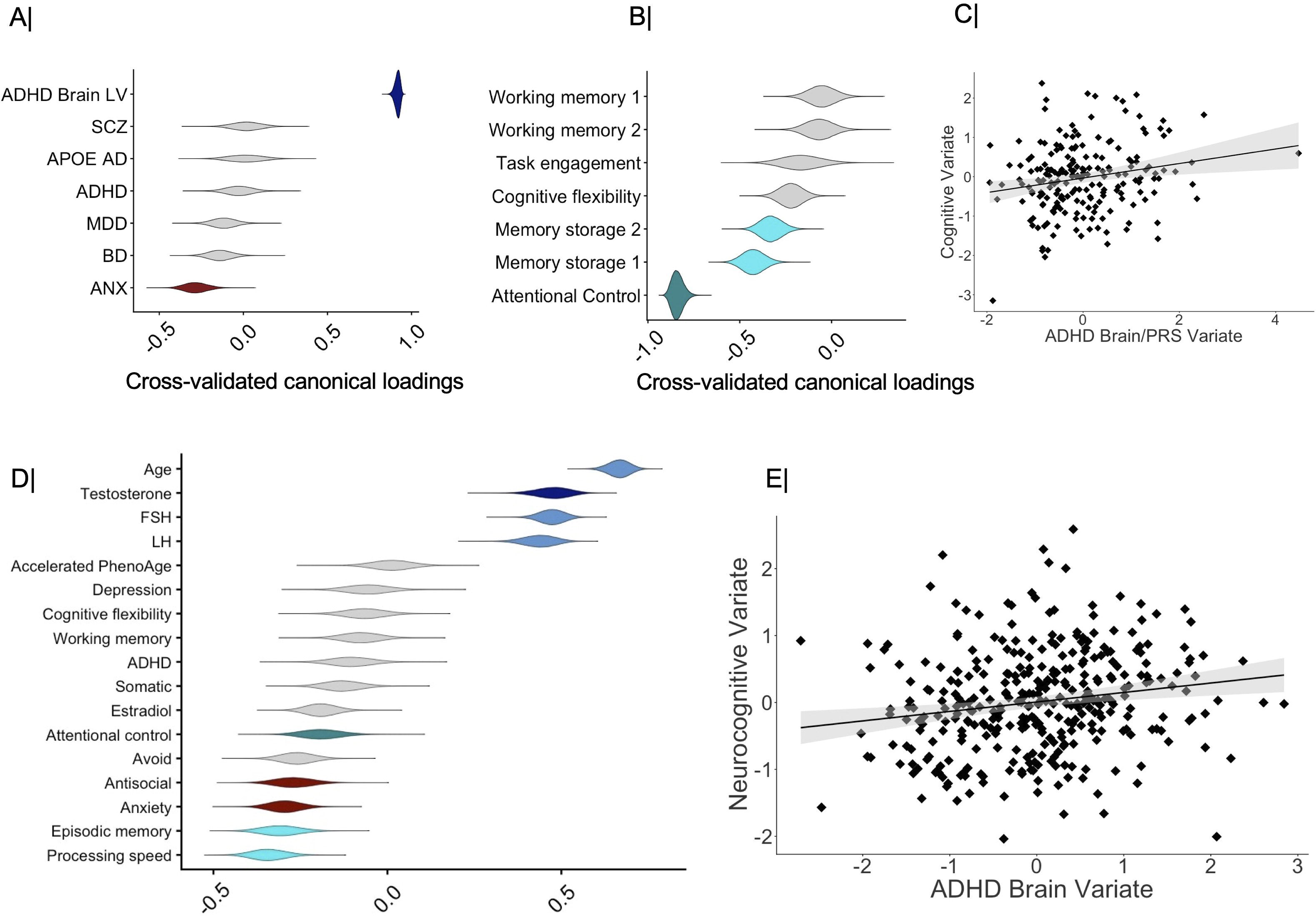
The brain LV from Analysis 2 and the PRSs associated with an ADHD diagnosis in Analysis 1 linked by CCA to cognitive performance and physiology in the Oregon (panels A- C) and HCP-A (panel D, E) samples. Panels A, B (Oregon) and D (HCP-A) show the partial correlation coefficients describing the relationship of the measured cognitive and polygenic risk (Oregon) or PhenoAge and hormone (HCP-A) scores with the predicted value of their corresponding canonical variate across all test CCA folds. The violin plots in panels A, B and D depict the distribution of partial correlation coefficients across the 100,000 bootstrap samples from the cross-validation procedure. Gray colored violin plots indicate non-robust canonical loadings (based on the bootstrapped 99% CIs from the cross-validation procedure). Panel C contains the scatter plot describing the linear relationship between the CCA variates pictured in panels A and B (Oregon). Panel E contains the scatter plot describing the linear relationship between the CCA variate pictured in panel D and the brain LV from Analysis 2 projected on rest and inhibitory control brain (HCP-A) data (see Results for details). PRS = polygenic risk score. CI = confidence interval. Task Engagement = accuracy in the zero-back condition (n-back task). Memory storage 1 = accuracy in the one-back condition (n-back task). Memory storage 2 = accuracy on the forward spatial span task. Working memory 1 = accuracy in the two-back condition (n-back task). Working memory 2 = accuracy on the backward spatial span task. LH = luteinizing hormone. FSH = follicle stimulating hormone. ADHD = attention deficit hyperactivity disorder. Antisocial = antisocial personality disorder. Avoid = avoidant personality disorder. Somatic = somatic disorder. Anx = anxiety. MDD = major depressive disorder. SCZ = schizophrenia; BD = bipolar disorder. AD = Alzheimer’s Disease.

#### Control analysis: Sex-dependent effects

To investigate whether preferential alignment with male-, rather than female-, typical FC patterns shows distinguishable relationships with ADHD-relevant cognitive challenges among boys vs girls, we re-ran Analysis 3, this time though introducing sex in the same CCA set as the cross-validated brain LV from Analysis 2 and the six PRSs robustly related to ADHD in Analysis 1. The resulting CCA mode was successfully cross-validated (*r*_CV_ = .26, 95% CI = [.13; .43], permutation-based *p* = 28 x 10^-5^) and replicated the canonical loading structure from Analysis 3 (Figure 4- A, B). Importantly, sex did not show a robust loading on its corresponding variate (loading_CV_ = .13, 95% CI [-.04; .31]]) which implies that the profile of relatively stronger alignment with male-typical FC, as determined in Analysis 2, shows equally strong associations with ADHD-relevant cognitive difficulties among boys and girls.

### 3.4 Analysis 4: The pattern of preferential alignment with male-typical brain architecture observed among children with ADHD correlates with gonadal hormone levels and poorer cognitive outcomes in healthy aging

Our fourth analysis tested whether patterns of alignment with female vs male-typical brain architecture linked to childhood ADHD across sexes would overlap with those related to greater cognitive decline and accelerated metabolic aging in older adulthood. To this end, we projected the brain LV from Analysis 2 onto the resting state and attentional control-related task data (i.e., Go/No-Go, (Bookheimer et al., 2019)) from the HCP-A sample.

Inclusion of the Go/No-go neuroimaging data allowed us to directly test the task relevance of the ADHD neural profile estimated in the Oregon study. Age was introduced in the CCA because we sought to determine whether individual differences in the expression of the ADHD brain profile would track with differences in cognition and circulating hormone levels which emerge with advancing years.

In the cross-validation of the estimated CCA model we controlled for the participant’s sex, scan site, average in-scanner motion during rest and task, race, handedness, education level, income-to-needs and regular medication use. The extracted CCA mode linked the ADHD profile of preferential alignment with male-typical FC during both rest (loading_CV_ of .56, 95% CI = [.44; .65]) and attentional control performance (loading_CV_ of .97, 95% CI = [.96; .98]) to higher testosterone, FSH and LH levels, poorer attentional control and episodic memory, slower processing speed, but reduced anxiety and antisocial personality symptoms among older individuals of either sex (Figure 5-C), *r*_CV_ = .17, 95% CI = [.06; .27], permutation-based *p* = .002 (Figure 4-D, E).

**Figure 5.**
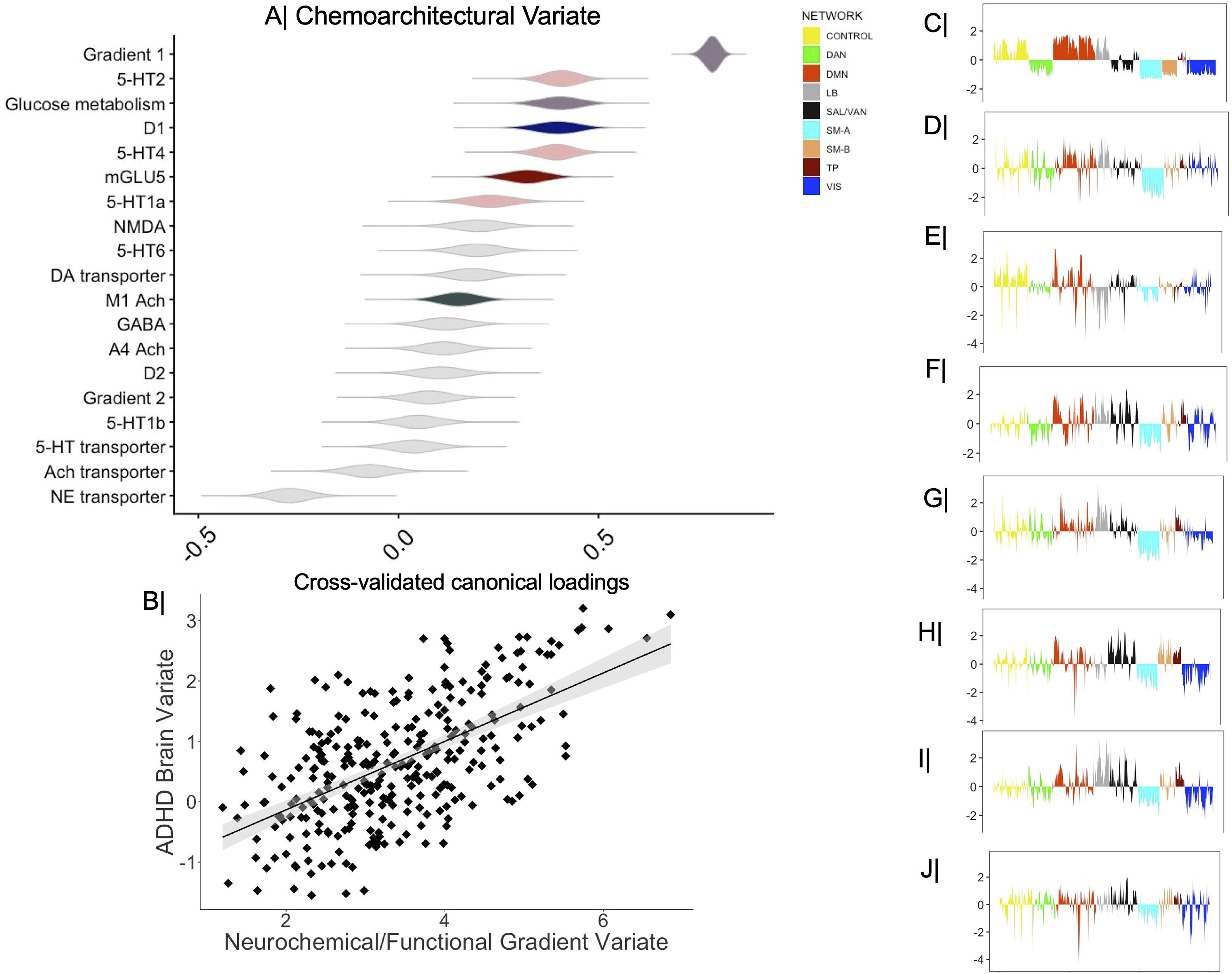
The functional gradient and neurotransmitter maps linked by CCA to the cross- validated brain LV from Analysis 2. Panel A shows the partial correlation coefficients describing the relationship of the functional gradient and receptor density maps with the predicted value of their corresponding canonical variate across all test CCA folds. The violin plots depict the distribution of partial correlation coefficients across the 100,000 bootstrap samples from the cross-validation procedure. In panel A, gray colored violin plots indicate non-robust canonical loadings (based on the bootstrapped 99% CIs from the cross-validation procedure). Panel B contains the scatter plot describing the linear relationship between the cross-validated brain LV from Analysis 2 and the CCA variate pictured in panel A. Panels C-J show the expression level/density map of the Gradient 1 (C), 5-HT2 (D), glucose metabolism (E), D1 (F), 5-HT4 (G), mGlu5 (H), 5-HT1a (I) and M1 Ach (J) aggregated by the Gordon networks. LV = latent variable. CI = confidence interval. 5-HT =serotonin. Ach = acetylcholine. D= dopamine. GABA = gamma-aminobutyric acid; GLU = glutamate. NMDA = N-methyl-D-aspartate receptor. NE = norepinephrine. Schaefer networks: TP= temporo- parietal. SAL-VAN = salience/ventral attention. LB = limbic. DMN = default mode. DAN = dorsal attention. SM-A = somatomotor-A. SM-B =somatomotor-B. VIS = visual.

### 3.5 Analysis 5: The ADHD-related alignment with male-typical FC emerges in areas of highest metabolic activity and dopaminergic (D1), serotonergic and glutamatergic receptor density

The results of our analyses so far raised the possibility that, across sexes, regional patterns of alignment with sex-typical brain architecture linked to childhood ADHD overlap with those related to steeper neurophysiological aging in older adulthood. To explore this possibility, we next investigated the spatial overlap between the ADHD brain map, characterized in Analysis 2, and the density map of receptors and transporters implicated in ADHD and AD (GABA/glutamate [GLU]), attention and learning (norepinephrine [NE], acetylcholine [Ach], dopamine [DA]), as well as mood and broad cognition (serotonin [5- HT]). To align our results with canonical representations of brain function, we also included the glucose metabolism map described in ref. (Castrillon et al., 2023) and the first two functional gradients identified by ref. (Margulies et al., 2016) which distinguish unimodal from transmodal (gradient 1 [FG1]) and somatomotor from visual (gradient 2 [FG2]) regions.

A CCA linked the ADHD brain map from Analysis 2 to the glucose metabolism, FG1, FG2 and receptor/transporter density maps listed above. Results indicated that the increased alignment with male-typical FC linked to childhood ADHD and steeper neurocognitive aging (Figure 5-A) is observed primarily in transmodal areas with greater metabolic activity and higher density of glutamatergic (mGLU5), serotonergic (5-HT1a, 5-HT2, 5-HT4), cholinergic (M1) and dopaminergic (D1) receptors, *r*_CV_ = .57, 95% CI = [.50; .66], *p_spin_* = 10^-5^ (Figure 5- B).

## 4. Discussion

Growing awareness that psychological well-being has not only immediate, but also temporally distant determinants, has fuelled commitment to a life course approach to healthy aging, which considers both individual-specific and structural contributors, including those that transcend generational boundaries (Hernandez et al., 2025; van Zwieten et al., 2025).

Heeding these calls for a more holistic understanding of mental health, we sought to characterize the potential neurobiological overlap between earlier life conditions with sex-biased prevalence and sex-related patterns of cognitive aging. Leveraging multimodal data from a child sample enriched for ADHD, we identified a profile of genetic risk which was independent of sex and psychiatric history. Specifically, in line with extensive reports on the heterogenous aetiology of this condition (Koirala et al., 2024; Lobo et al., 2025), we found that boys and girls diagnosed with ADHD were at higher genetic risk not only for ADHD, but also, albeit lesser extent, for MDD, ANX, SCZ, BD and APOE-related AD. The near-zero correlation of the APOE-related AD PRS with the remaining PRSs underscores its potentially unique relevance to the clinical presentation of ADHD, thereby raising the possibility that neurodevelopmental disorders could help shed light on the subclinical markers of suboptimal aging trajectories.

The pathophysiology of ADHD is allegedly best captured by holistic, rather than localizationist, brain approaches (Bedford et al., 2025; Koirala et al., 2024). Accordingly, our analyses linked increased whole-brain similarity to male, rather than female architecture to ADHD, including its typical differences in inhibitory control and memory storage (Koirala et al., 2024; Thapar & Cooper, 2016). In other words, a diagnosis of ADHD was observed among boys who showed hyper-alignment with male-normative FC patterns and girls who deviated from female-normative FC patterns. Speaking to the feasibility of identifying early life biomarkers of suboptimal lifespan developmental trajectories, our analysis of the HCP-A sample related the neural profile of childhood ADHD to markers of reproductive aging (i.e., increased FSH and LH levels (Ahamed et al., 2026; Bhatta et al., 2018)), as well as differential performance deficits on ADHD- (i.e., attentional control) and aging-relevant (i.e., episodic memory, processing speed) tasks (Frisoni et al., 2022) among older adults of either sex. Testifying to its likely functional involvement, projection of the ADHD profile onto the resting state and the inhibitory control-related data from the HCP-A sample yielded stronger associations with aging in the task data. In line with the key role of gonadal hormones in neural sexual differentiation (Gegenhuber et al., 2022; Küchenhoff et al., 2024), in the HCP- A sample, stronger expression of the ADHD brain profile was detected among older men and women who showed relatively higher levels of testosterone. Since testosterone declines with age across both sexes (Ahamed et al., 2026), the observed effect, suggestive of maintenance, rather than decline, likely speaks to how individual differences in gonadal hormone levels may shape regional variability in alignment with sex-typical brain architecture and, thus, indirectly mental health outcomes. Unavailability of hormonal measures in the Oregon sample precluded more in-depth tests of this hypothesis, which we believe warrants further study.

ADHD-related alignment with male-typical FC was most salient in association (rather than unimodal) regions (Margulies et al., 2016), particularly those showing greater metabolic activity (Castrillon et al., 2023) and higher density of glutamatergic (mGLUR5, NMDA), muscarinic, dopaminergic (D1) and serotonergic (5-HT_2A,_ 5-HT_4_) receptors. These results dovetail with evidence implicating the association networks, specifically, the DMN and its connections with control and attentional areas, in the pathophysiology and inhibitory control challenges connected to ADHD (Anderson et al., 2025; Koirala et al., 2024) and pathological aging conditions such as AD (Kvavilashvili et al., 2020). The spatial overlap between the 5- HT_2_/ 5-HT_4_ receptor density maps and the brain profile of ADHD linked to higher MDD, ANX and BD PRS reaffirms the role of 5-HT in mood pathology (Erritzoe et al., 2023). The relevance of the cholinergic and dopaminergic (D1 receptors) systems to childhood ADHD and neurocognitive aging reasserts the importance of these neurotransmitters to working memory and broader cognitive control processes across the life course (Ananth et al., 2023; Ballinger et al., 2016; Ciampa et al., 2022; Gustavsson et al., 2023; Joyce et al., 2025), particularly online maintenance of mental representations for D1 receptors (Braun et al., 2021; Durstewitz & Seamans, 2008), all of which are deficient in ADHD (Koirala et al., 2024) and pathological aging conditions such as AD (Frisoni et al., 2022; Hampel et al., 2018).

Also noteworthy is the spatial overlap between glutamatergic receptors, specifically mGLUR5, and the brain profile linked to ADHD and neurocognitive aging. This correlation not only reaffirms the importance of GLU dysregulation to ADHD (Vidor et al., 2022), but also resonates with cross-species evidence on the role of GLU in the pathophysiology of dementia linked to AD (Dupuis et al., 2023), particularly the involvement of mGLUR5 signalling in early AD (Mecca et al., 2020; Wang et al., 2024).

Complementing the reported connection between mood problems and stronger alignment with female-, rather than male--, typical FC patterns along the transmodal-to-unimodal functional hierarchy in adolescence (Petrican et al., 2025)), anxiety disorder scores correlated negatively with the ADHD profile of alignment with male-typical FC among the older HCP-A adults. Unexpectedly, antisocial personality scores showed a negative association with the ADHD brain profile, thereby raising the possibility of age-dependent associations between alignment with sex-typical brain architecture and distinct externalizing spectrum conditions.

Our investigation has several limitations. First, our analyses featured predominantly white youths and adults. Given the under-recognition and under-diagnosis of ADHD in minoritized groups (Madsen et al., 2018; Mennies et al., 2021; Prasad et al., 2019; Wright et al., 2015), larger and more demographically diverse samples would help extend the effects herein reported, particularly by allowing sub-typing of the ADHD and aging groups. Second, we used sex-independent PRSs. More extensive development and validation of sex-specific markers would facilitate a more fine-grained investigation of the neurobiological links between childhood ADHD and aging outcomes. Third, due to data availability, hormonal and physiological aging were assessed only in the adult sample, the latter via an organism-level, sex-independent measure (Levine, 2013). Collection of multiple markers of “wear-and-tear”, including stress hormone levels, pace of aging based on blood chemistry data (Balachandran et al., 2025), correlated patterns of physiological and epigenetic senescence (Levine et al., 2018), specific inflammatory processes (innate vs adaptive) in different tissue and cell types (Grandke et al., 2025; Zhang et al., 2022), would allow a more detailed comparison of the multiscale biological processes likely to connect neurodevelopmental disorders to neurodegenerative risk.

In summary, childhood ADHD was linked to higher genetic risk for multiple psychiatric conditions beyond ADHD (MDD, ANX, SCZ, BD) with an independent association observed for APOE-related AD. Boys and girls diagnosed with ADHD showed preferential alignment with male-typical, rather than female-typical, FC in transmodal areas with the highest dopaminergic (D1), serotonergic, glutamatergic and muscarinic receptor density. In a separate sample, the brain profile of childhood ADHD correlated with higher levels of testosterone, hormonal markers of reproductive aging (i.e., increased FSH and LH levels) and poorer performance on cognitive tasks relevant to pathological aging among older individuals of either sex. Our results underscore the feasibility of identifying multiscale biomarkers linking neurodevelopmental conditions to suboptimal adult developmental outcomes, which could enable timely and more effective psychiatric interventions.

## Materials & Correspondence

Correspondence and material requests should be addressed to R.P.

## Data statement

The raw data are available at https://nda.nih.gov/study.html?id=1938 (DOI: 10.15154/1528485) (Oregon), at https://db.humanconnectome.org (HCP) and at https://nda.nih.gov/ccf (HCP-A) upon completion of the relevant data use agreements. The HCP-A data used in this report came from Data Release 2.0 (DOI: <u>10.15154/krqa-9963</u> ).

## Code availability

We used already existing code, as specified in the main text with links for free download.

## Acknowledgments

Data used in the preparation of this article were obtained from the Oregon ADHD-1000: A longitudinal data resource enriched for clinical cases and multiple levels of analysis, Human Connectome Project, WU-Minn Consortium (Principal Investigators: David Van Essen and Kamil Ugurbil; 1U54MH091657; funders: the 16 NIH Institutes and Centers that support the NIH Blueprint for Neuroscience Research and the McDonnell Center for Systems Neuroscience at Washington University), as well as from the Human Connectome Project-Aging (HCP-A) Study (https://humanconnectome.org/study/hcp-lifespan-aging). The Oregon ADHD-1000 project was supported by multiple NIMH grants: 1R01-MH59105 (Nigg); R01-MH63146 (Nigg); R01-MH070004 (Friderici); 1R01MH86654 (Nigg-Fair); R01MH099064 (Nigg); 1R37MH59105 (Nigg); 2R56MH086654 (Nigg); R01MH115357 (Fair-Nigg); 2R37MH59105 (Nigg); R01 MH096773 (Fair); K23 MH108656 (Karalunas). Data storage at OHSU is supported by the Oregon Clinical and Translational Research Institute funded by a grant from the National Center for Advancing Translational Sciences (NCATS), National Institutes of Health, through Grant Award Number UL1TR002369.

The HCP-A study is part of the HCP-Lifespan research project, which is supported by grants U01MH109589, U01MH109589-S1, U01AG052564, and U01AG052564-S1 and by the 14 NIH Institutes and Centers that support the NIH Blueprint for Neuroscience Research, by the McDonnell Center for Systems Neuroscience at Washington University, by the Office of the Provost at Washington University, by the University of Minnesota Medical School, by the University of Massachusetts Medical School, and by the University of California Los Angeles Medical School. This manuscript reflects the views of the authors and may not reflect the opinions or views of the NIH, HCP, or Oregon ADHD 1000 consortium investigators.

